# RT States: systematic annotation of the human genome using cell type-specific replication timing programs

**DOI:** 10.1101/394601

**Authors:** Axel Poulet, Ben Li, Tristan Dubos, Juan Carlos Rivera-Mulia, David M. Gilbert, Zhaohui S Qin

## Abstract

The replication timing (RT) program has been linked to many key biological processes including cell fate commitment, 3D chromatin organization and transcription regulation. Significant technology progress now allows to characterize the RT program in the entire human genome in a high-throughput and high-resolution fashion. These experiments suggest that RT changes dynamically during development in coordination with gene activity. Since RT is such a fundamental biological process, we believe that an effective quantitative profile of the local RT program from a diverse set of cell types in various developmental stages and lineages can provide crucial biological insights for a genomic locus. In the present study, we explored recurrent and spatially coherent combinatorial profiles from 42 RT programs collected from multiple lineages at diverse differentiation states. We found that a Hidden Markov Model with 15 hidden states provide a good model to describe these genome-wide RT profiling data. Each of the hidden state represents a unique combination of RT profiles across different cell types which we refer to as “RT states”. To understand the biological properties of these RT states, we inspected their relationship with chromatin states, gene expression, functional annotation and 3D chromosomal organization. We found that the newly defined RT states possess interesting genome-wide functional properties that add complementary information to the existing annotation of the human genome.

**AUTHOR SUMMARY:** The replication timing (RT) program is an important cellular mechanism and has been linked to many key biological processes including cell fate commitment, 3D chromatin organization and transcription regulation. Significant technology progress now allows us to characterize the RT program in the entire human genome. Results from these experiments suggest that RT changes dynamically across different developmental stages. Since RT is such a fundamental biological process, we believe that the local RT program from a diverse set of cell types in various developmental stages can provide crucial biological insights for a genomic locus. In the present study, we explored combinatorial profiles from 42 RT programs collected from multiple lineages at diverse differentiation states. We developed a statistical model consist of 15 “RT states” to describe these genome-wide RT profiling data. To understand the biological properties of these RT states, we inspected the relationship between RT states and other types of functional annotations of the genome. We found that the newly defined RT states possess interesting genome-wide functional properties that add complementary information to the existing annotation of the human genome.

## INTRODUCTION

The functional significance of genome organization at the sequence level of many regions in the human genome remains poorly understood despite the completion of the human genome sequencing project more than 15 years ago. Large international consortia such as ENCODE (1), REMC (2) and IHEC (3) have demonstrated that there are many functional elements present throughout the genome that play interesting and important roles in structuring the human genome. The discovery of these functional elements highlights the importance of defining, cataloguing, and characterizing the diverse regions that remain poorly understood. It is important to advance the field through a variety of methods, integrating different experimental designs and findings, allowing the definition of structure like topologically associating domain (TADs) (4-6) or Lamin Associated Domains (LADs) (7, 8). Alternatively, statistical modelling applied on these experimental data created a new, composite types of genomic elements. As a prime example, Ernst and Kellis exploited ChIP-seq data of histone modifications to classify the genome into chromatin states using the Hidden Markov Model (HMM) (9-11). They found that many of these chromatin states are correlated with particular cis-regulatory elements, such as enhancers or insulators. The appealing feature of the chromatin state concept is its ability to crystallize complex and biologically meaningful patterns into a few numbers using a large collection of epigenomic profiles that contain lots of variation, overlap and redundancy. We believe the same strategy can be applied to other types of data. In this study, we annotate the genome using a rich set of genome-wide replication timing (RT) programs derived from multiple human cell types (12, 13). The RT programs represent the genome duplication occurring in a defined temporal order in human cells.

In humans, the RT program is dynamic and cell type-specific. The difference between types of differentiated cells, and even stem cells and their progenitors can be dramatic. For example, half of the genome undergoes RT changes during cell differentiation of embryonic stem cells (14, 15). These dynamic changes in RT occur in units of 400-800 kb known as replication domains (RDs) (12, 16, 17). RT alterations have also been linked to many diseases (18-21). Recent studies have revealed an association between RT and higher-order chromosomal structure (TADs and sub-nuclear compartment A and B) (17, 22-26), as well as gene expression regulation (24), supporting the hypothesis that the genome is partitioned into smaller replication units of replication (27). In general, early replication is correlated with elevated transcriptional activity (17, 28). Researchers have classified RDs as constitutive or developmentally regulated, depending on how consistent the replication occurs across different cell types (17, 28). RD boundaries are stable across different cell types, but the changes of RT on the developmental domains are correlated with changes in sub-nuclear position, chromatin structure and transcriptional state.

The introduction of RDs helps put context to a genomic locus from the RT perspective, but the concept is qualitative. As more RT profiles of more cell types becomes available, we gain the opportunity to characterize RT dynamics in a more refined and quantitative fashion. In this study, we use 42 RT profiles collected from 25 cell types and differentiation intermediates (24), and apply a Hidden Markov Model (HMM) based approach to break down the genome into 15 RT states. The goals are two-fold: first, identify frequent combinatorial RT patterns among a compendium of cell types; second, annotate the genome by assigning an RT state to every locus. Note that unlike chromatin states (9-11), RT states defined here remain unchanged across cell types since they represent the behaviour of chromatin across a variety of cell types, rather than the composition of chromatin in a particular cell type. The 25 cell types and the differentiation intermediates have been strategically selected to represent the diversity among all the cell types in human and represent developmental stages of the three germ layers (24). Finally, the RT states described here represent a more accurate and quantitative means of defining constitutive early, constitutive late or developmental RDs, which adds a novel quantitative annotation of the functional properties of the human genome.

## RESULTS

### RT value distribution along the genome

RT values from Repli-chip (12, 13) are the log2 ratios of signals derived from early and late samples measured by microarrays from published studies (24). About 85% of the genome is covered by RT values. The dynamic range has been normalized which ranges from-2.9 to 2.9 as previously described (24, 29), and their distributions in all cell types show a similar bi-modal pattern with two peaks around-1 and 1, representing respectively the genome part replicate late and early during replication (Fig 1A). Higher RT values (>0.3) were classified as early replicating; low values (<-0.3) were classified as late replicating (24). Alignment of RT values from 42 data sets along a chromosome shows the global preservation of RT during development and also some deviations among cell types (Fig 1B). As examples, we show some regions with early RT values in hESC cells but late in other cell types (the green rectangle in Fig 1B). The opposite pattern can be observed in the blue rectangle (Fig 1B). Such patterns illustrate the importance of statistical modelling of RT program to identify recurrent and biologically meaningful combinatorial patterns of RT values.

**Fig 1:**
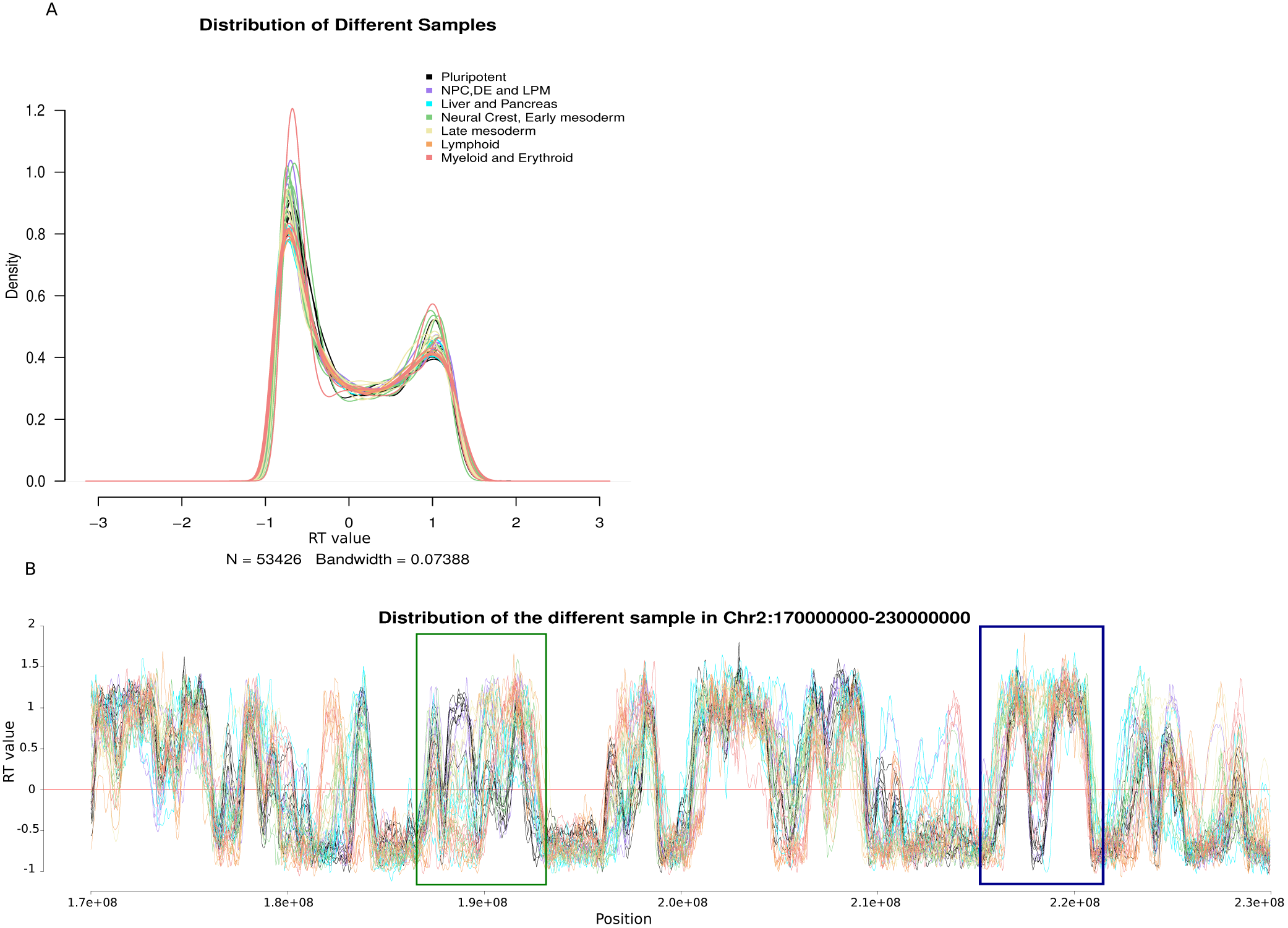
Properties of RT values. **A.** Global RT value distributions along the 42 datasets of 25 cell types. **B.** Aligned tracks of RT values in the chr2:170,000,000-230,000,000 region across all samples. The y axe is the RT value between-1.5 and 1.5 in this region; the x axe is the genome coordinate. The red dash line show the RT value of zero. The rectangles highlighted variations among cell lines. In the green box we observe a region with early RT value in pluripotent and MED cell lines but late or close to zero RT values in Lymphoid, Myeloid and Erythroid cell lines. In the blue box we observe a region that shows early RT values in Mesoderm, Liver and Pancreas cell lines but late RT values in pluripotent and MED cell lines. The same key color is used in A and B panel.

### Genome decomposition into RT states

The HMM requires the specification of the number of hidden states. In this study, we chose *K*= 15 to balance complexity and biological interpretation. We have tested several different numbers of states (6, 10, 20, 25 and 30) and found the results are largely comparable (Supplementary Figure 1 to 5). In addition, we found that using 15 states, clustered the cell lines according to their genome-wide RT profiles in a manner consistent with the lineage origin of each cell type (Fig 2) (24). We analysed the spatial organisation of the RT states, i.e., how often does the switch (RT transition of contiguous regions) between different RT states occur. By looking at the RT changes in neighbouring genomic regions, we found that the RT states tend to stay unchanged for long stretches of chromosomes. It is evident from the transition matrix that the probability of staying in the same state is very high and immediately adjacent chromosomal segments are mostly classified into the same RT state (Supplementary Figure 6A). Analysis of the average of contiguous bins for each RT states shows that the two largest domains are observed for the Early1 (618 kb) and Late3 (1.16 Mb). The other RT states have value between 267 kb and 500 kb (Supplementary Figure 6D). This is consistent with the previous findings that RDs are usually hundreds of kilobases in length (12, 16, 17).

**Fig 2:**
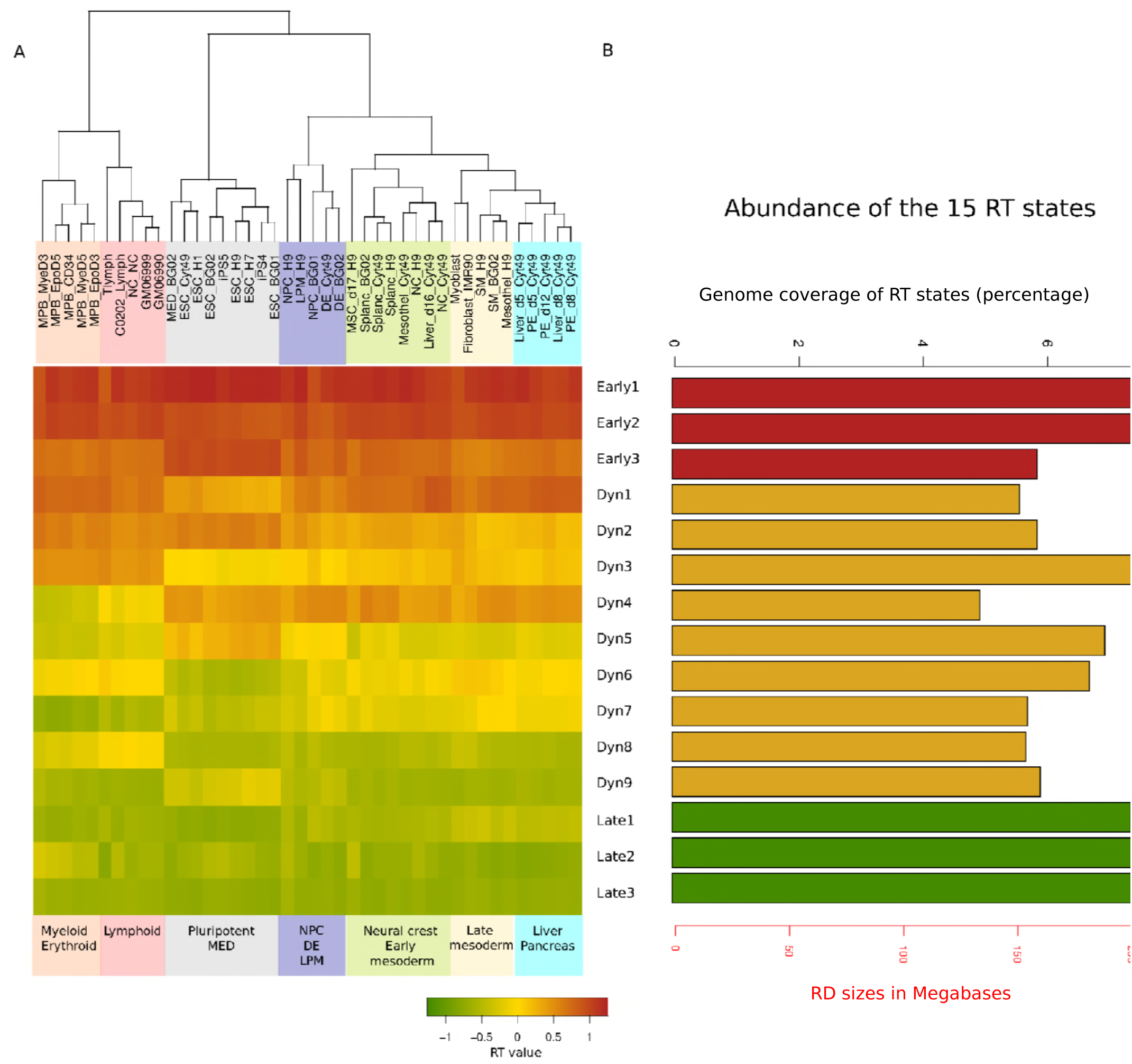
Overview of 15 RT states. **A.** RT values of the 15 RT states along the 42 datasets. The genome was decomposed into 15 RT states based on the RT program of each sample. The average value for each state is displayed in the heatmap and the scale is displayed at the bottom. The dendrogram was produced by hierarchical clustering of the samples. **B.** Percentage and size (in mega bases, red axis) of genome coverage for each of the 15 RT states.

### Overview of the 15 RT states

From Fig 2A, we saw that the RT states Early1 and Early2 replicate early across almost all cell types; Early3 shows significantly earlier RT in pluripotent cell types than in differentiated cell types. The RT states which change between cells are called dynamic RT state (Dyn). RT state Dyn1 displays significantly earlier RT in differentiated cell types. For RT state Dyn2, pluripotent cells and myeloid/erythroid and lymphoid cells show significantly earlier RT than other cell types. In RT state Dyn3, pluripotent cells show later RT than other cell types. In RT state Dyn4, myeloid/erythroid and lymphoid cells show later RT than other cell types. RT state Dyn5 shows a trend similar to the Early3 RT state, with intermediate RT in pluripotent cells but significantly later RT in all other cell types. RT Dyn6 shows late RT in pluripotent cells and intermediate RT for the rest. RT state Dyn7 shows late RT among myeloid/erythroid and lymphoid cell types and intermediate RT elsewhere. RT state Dyn8 shows the opposite pattern in Dyn8, i.e., intermediate RT among myeloid/erythroid and lymphoid cells but late RT for the rest. RT state Dyn9 shows the opposite pattern in Dyn6, i.e., intermediate RT in pluripotent cells but late RT elsewhere. RT states Late1-3 show late RT across all cell types in general. In Late1, late mesoderm and liver/pancreas cells tend to show less late RT compared to other cell types, whereas Late2 shows less late RT in myeloid/erythroid cell types compared to other cell types.

### Global RT organisation of the defined states

The distribution of the 15 RT states across the genome is shown in Fig 2B. Since we excluded centromeric regions, approximately 85% of the genome is covered by RT state assignment. The proportion of the genome represented by each state is similar across all RT states, ranging from 132 Mb (4.9% genome coverage, RT state Dyn4) to 217 Mb (8.1%, RT state Early1), with a mean of 168 Mb.

We also analysed the distribution of RT states based on the gene density per chromosome (Fig 3). The distribution reveals that high gene-density (HGD) chromosomes (chr1, 11, 12, 15, 16, 17, 19 and 22) contain regions with more early RT states and low gene-density (LGD) chromosomes (chr4, 6, 8, 13, 18, 21) contain regions with more late RT states (also see Supplementary Figure 6B and C) (30). We then analysed the base composition of the defined RT state and found a monotonically increasing trend in AT percentage from early RT states to late RT states: Early1 RT state has around 52% AT, compared to about 64% AT in the Late3 RT state (Supplementary Figure 6E). This is expected since most of the late RT states are located in gene-poor regions, which are marked with high AT content (31).

**Fig 3:**
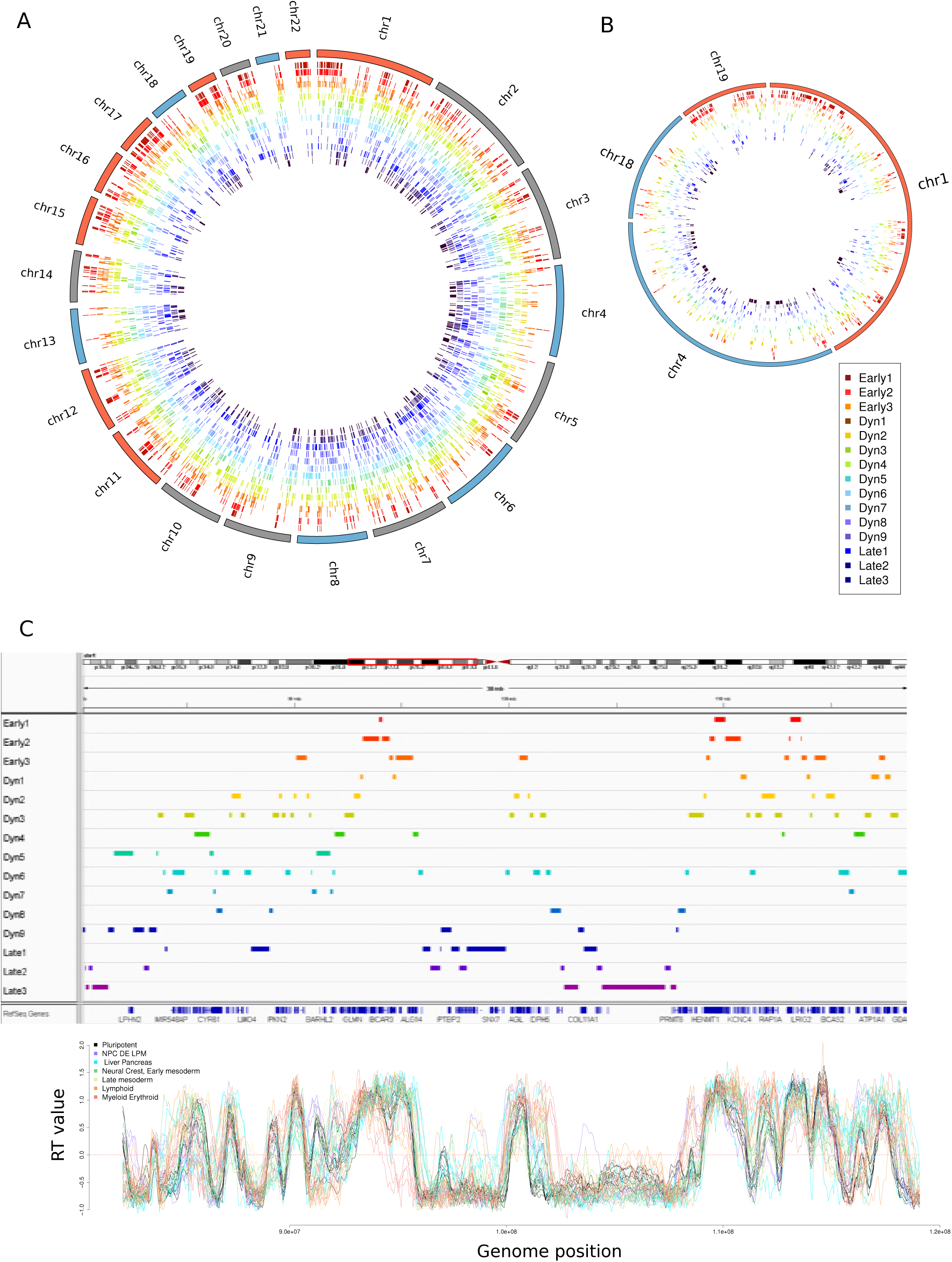
Distribution of the 15 RT states along the genome. High gene density (HGD) chromosomes are highlighted by outside rectangles in red, low gene density (LGD) chromosomes are highlighted by outside rectangles in blue and all the other are in represented in grey. **A.** Circos plot shows the whole genome distribution. **B.** Circos plot of RT states distribution for selected HGD chromosomes (chr1 and 19) and LGD chromosomes (chr4 and 18). HGD chromosomes have larger and dense Early domains and smaller and sparse Late domains while LGD chromosomes have an opposite pattern. **C.** IGV screenshot of the 15 RT states distribution and also the raw replication timing along a 40 Mb segment of the chromosome 1.

### Periodic pattern of RT states observed in part of the genome

Illustrated by the RT states shown on a segment of chromosome 2 (Fig 3C), we observed an interesting wave-like pattern with RT state changing from early to late, then late to early in an ordered and recurrent fashion. The patterns are very clear and pronounced in about half of the genome and are chromosome-specific (Fig 3A and 3B). To study such wave-like pattern along the genome, we merged regions corresponding to the two earliest RT states (Early1 and Early2) and the two latest RT states (Late2 and Late3) into two different subsets called respectively Early domains and Late domains. We then computed for each subset (Early or Late) the distance between neighbouring domains. Due to the large difference of Early and Late domains distribution between chromosomes (Supplementary Figure 6B), we considered HGD chromosomes and LGD chromosomes separately. We found that the average size of Late domain in HGD chromosomes is 643 kb and the average distance between two neighbouring domains is 4.27 Mb. For LGD chromosomes, the average size of Late domain is 795 kb and 3.83 Mb for the average distance between two adjacent domains. For the Early domains in HGD chromosomes we found an average size of 894 kb and an average distance of 3.28 Mb between two neighbouring domains. In LGD chromosomes, the average size is 638 kb and the average distance between two adjacent ones is 5.88 Mb (results for each chromosomes presented in Supplementary Table 2 and 3). The analysis of the Early domain shows a significant difference between HGD and LGD in their sizes (p-value = 6.02E-04) and distances (p-value= 1.01E-02). But neither size nor distance in the Late domains showed significant differences (Supplementary Table 4).

RT states clearly illustrate that replication is consistently executed in an orderly fashion. And the RT profiles show distinct pattern between HGD and LGD chromosomes. Perhaps the HGD chromosomes replicated faster than the LGD chromosomes since the latter ones are less accessible. The imbalance of the Early and Late domains on the HGD and LGD chromosomes are similar to that of the A and B compartments discovered on the Hi-C experimental data (6, 32, 33). We speculate that the distinctive and elegant periodic pattern we observed in RT indicates the chromosomal properties are highly regulated and the pattern may contribute to its ability to be readily folded in 3D space in an ordered fashion (34).

### Annotation features of distinct RT states

To characterize the different RT states identified by our HMM method, we compared them to the previously described chromatin states annotated based on epigenetic modifications derived from ChIP-seq data (9). Additionally, we also analysed the gene expression activity inside different RT states.

First, we investigated the abundance of the four types of gene families (protein coding, ncRNA, lncRNA and pseudogenes) in each of the 15 RT states. The results are summarized in Fig 4. We saw that the proportion of bins with annotation is high in Early states and decreased steadily when moving from Early states to Late states. Only about 50% of the Late state bins possess at least one gene annotation (of four types), down from 98% in Early states (Fig 4A). In all RT states, protein-coding genes are the most frequently detected gene annotations except for the Late3 RT state where around 40% of the gene annotations are lncRNAs and only 35% are protein-coding genes. We found protein-coding gene annotation frequency decreased steadily across RT states from Early (60%) to Late (35%) (Fig 4B). Frequency of ncRNA annotations is stable throughout all the states (Fig 4C) and lncRNA frequency increased steadily from Early1 (≈ 15%) to Late3 (≈ 35%) (Fig 4D). Pseudogenes annotation is similar in Early and Dyn1-7 states and higher in the other states (Fig 4E). From the above, we concluded that early RT is positive correlated with the density of protein-coding genes, negatively correlated with the density of lncRNAs, but not correlated with the density of ncRNAs and pseudogenes.

**Fig 4:**
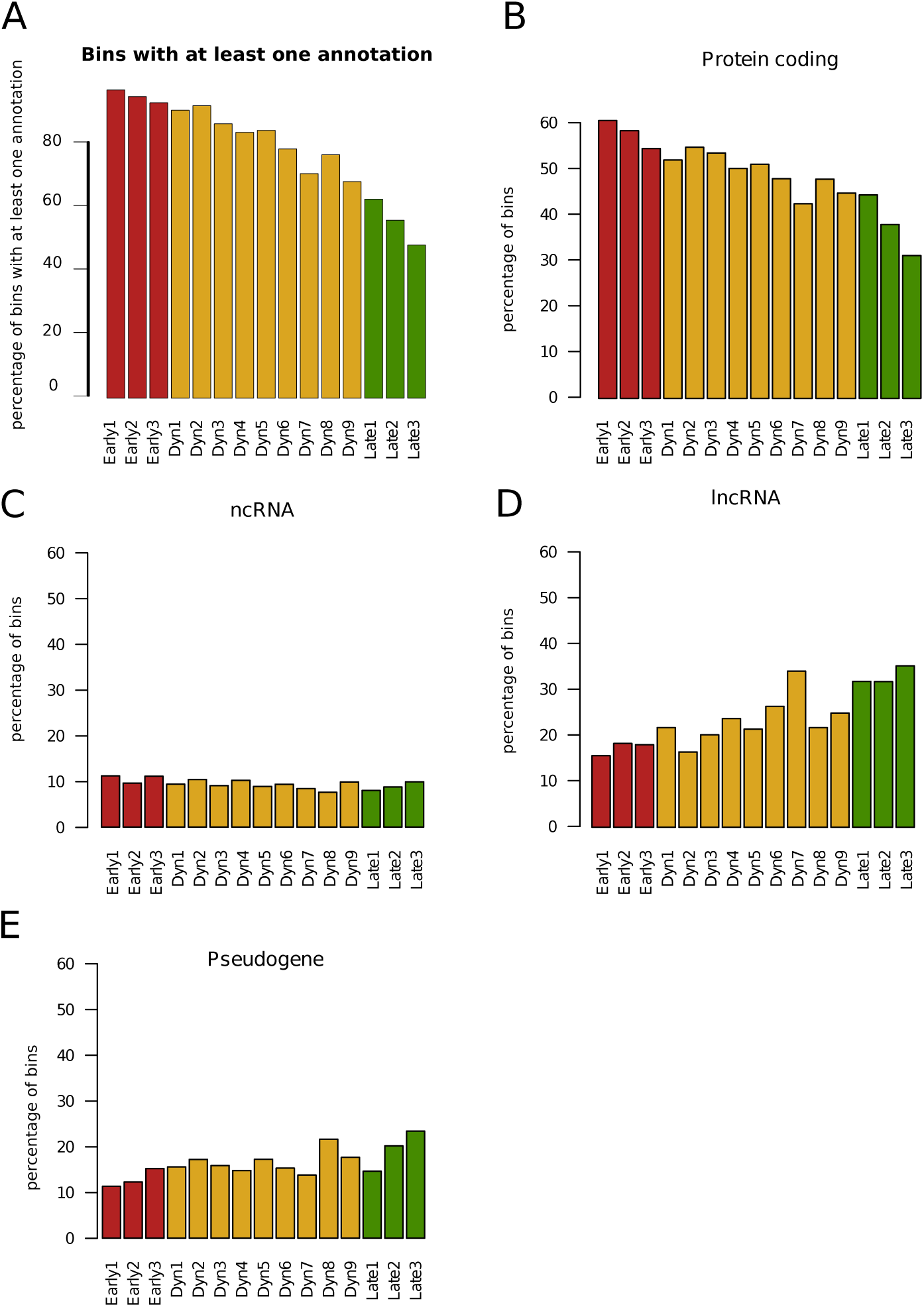
Annotation features of the different RT states. **A.** Percentage of bins with at least one of the following annotation: protein-coding, ncRNA, lncRNA or pseudogene. **B.** Protein coding annotation percentage by RT states. **C.** ncRNA annotation percentage by RT states. **D.** lncRNA annotation percentage by RT states. **E**. Pseudogene annotation percentage by RT states.

### Specific chromatin architecture is detected by combining RT states and chromatin states

It is of interest to compare the relationship between previously defined chromatin states (9) and newly defined RT states. Since chromatin states are cell type specific, we hope to show how the dynamic DNA chromatin organisation changes with respect to the DNA replication profiles in different cell lines. Fig 5A shows the average proportion of the genome belonging to the 225 different combinations of the 15 chromatin states (9) and the 15 RT states defined in this work. Overall, we saw that except for chromatin states 13-15, the proportions of all the other 12 chromatin states decrease steadily in regions corresponding to RT states moving from Early to Late. For chromatin state 13 (Heterochromatin), the pattern is reversed as expected: the proportions keep increasing with RT states moving from Early to Late. Interestingly, we found chromatin states 14 and 15 (repetitive/CNV1 and 2) are significantly enriched (chromatin state 14: 35% and chromatin state 15: 34%) in regions corresponding to RT state Dyn8. We also observed that these two CNV states have high proportions in Dyn9 and Dyn7. The CNVs have been shown to be correlated with later RT value and located in AT rich region (35). These are consistent with the characteristics of these three dynamic RT states which have an AT content around 60% (Supplementary Figure 6E) and relatively late RT. With the combination the RT states and the chromHMM states (Supplementary Figure 7), we observed that leukemia cell line K562 possesses a unique pattern with relatively higher proportion of CNVs chromatin state at the early RT states than the other cell lines. Again, we saw that the Dyn7, 8 and 9 RT states are enriched in CNV states in all cell lines (Supplementary Figure 7). In the RPKM analysis we also observed decreased RPKM values in Dyn3 RT state in the H1 cells in comparison of the cell lines used in the chromHMM analysis (Fig 5B). These two examples show the dynamics among RT, chromatin organisation and gene expression in different cell types.

**Fig 5:**
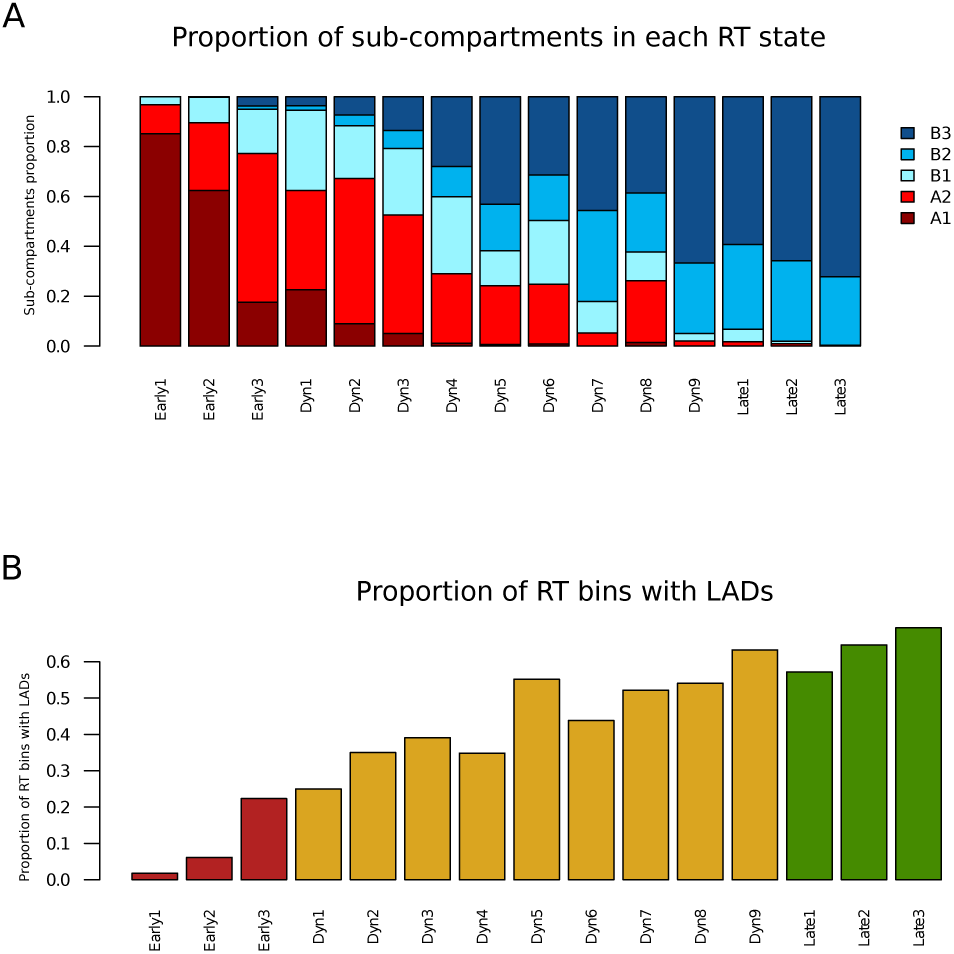
ChromHMM and transcription level analysis. **A**. shows the average value of the states by cell lines. Average of the ChromHMM states proportion by RT states of nine cell lines are displayed on the right of the figure (see material and methods). **B**. Expression analysis of protein coding genes along the 15 RT states in different cell lines (log10(RPKM) value is shown).

**Fig 6:**
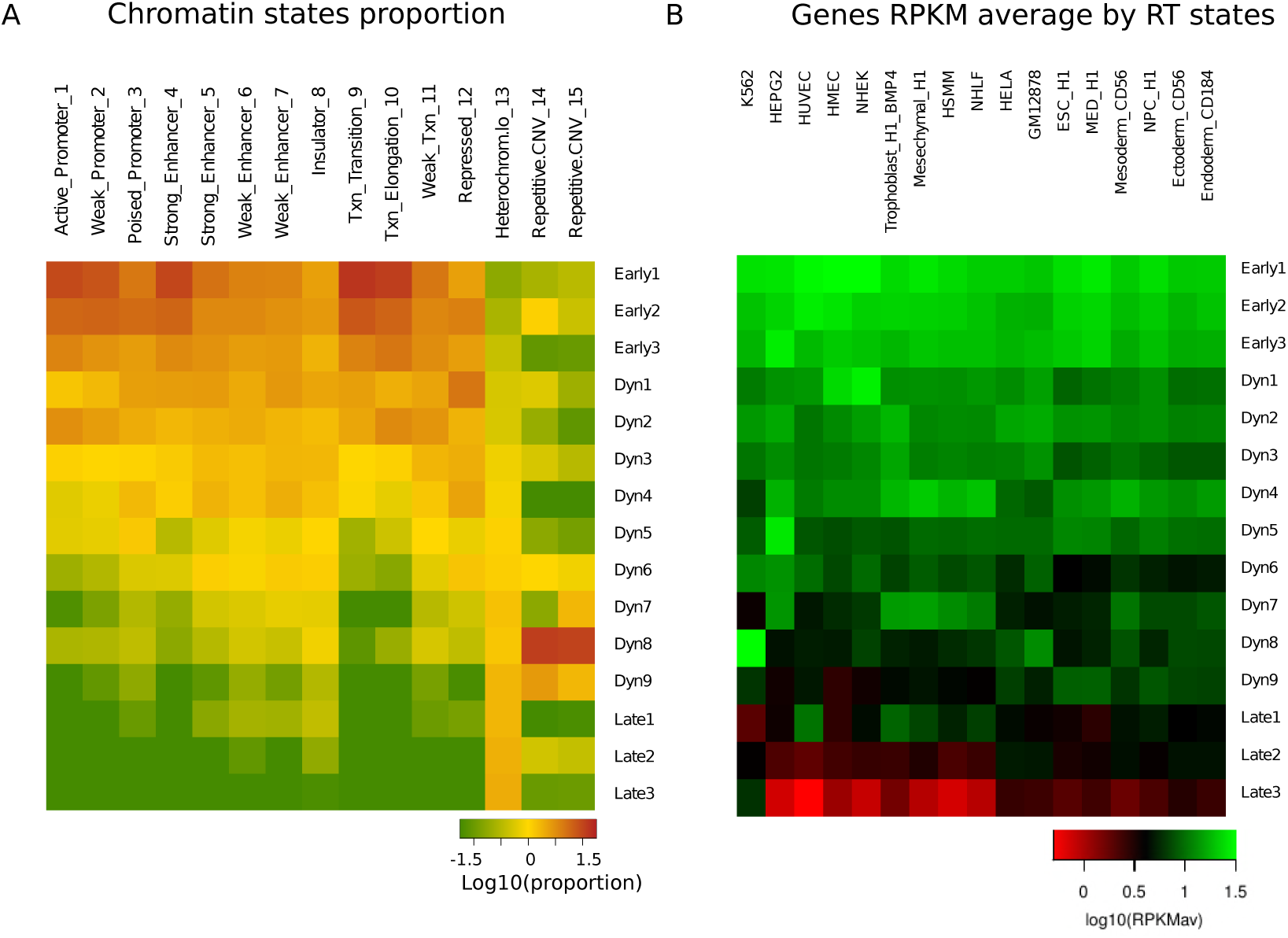
Comparison between RT states and 3D organisation of the genome. **A.** Composition of the five sub-compartments in each of the 15 RT states. **B**. Proportions of RT bins possessing LADs for each RT state.

The above observation suggests that the chromatin states and RT states contain complementary information. Either one of them alone is not enough to capture all the diversities observed in different epigenomes. As an example, regions annotated by the same chromatin state may show dramatic different RT state annotation and vice versa. Thus adding RT states on top of chromatin states enable a better annotation of the genome.

### Gene expression and GO term enrichment analysis in the RT states

For each cell line, we computed the average gene expression (in RPKM) for each RT state (Fig 5B). We first observed that the cell lines of the same type tend to cluster together. As expected, protein-coding genes located inside regions belonging to the three Early states are highly expressed. Genes located in genomic regions belonging to the Late3 RT state have the lowest expression values in all cell lines tested. Dynamic states are more complex and variable, with intermediate expression values between Early and Late states. These observations confirmed that regions with higher RT values tend to have higher gene expression levels.

To find out if there is any specific pathway or gene set that is enriched in different RT domains, we conducted a GO term enrichment analysis on the genes overlapping with each of the 15 RT states. The 20 most significantly enriched GO terms for each RT state are presented in Supplementary Table 5. We also calculated the average RT state score (since RT state number is ordinal) for each GO term and report result in Supplementary Table 5. We observed that Early states are enriched in processes involved in housekeeping pathways, like cell division, apoptosis, and transcription regulation (Table 1 and 2 and Supplementary Table 5). That is consistent with the RT score analysis which showed lower values for the same pathways (apoptotic process (3.99), protein phosphorylation (3.74), cell division (3.7)) (Supplementary table 5). Late RT states are enriched with GO terms about stress response or defence response, and are enriched in processes involved in neural cell regulation, and tissue development. These pathways possess high average RT scores (> 9) (Supplementary Table 5). Dyn RT states are enriched in transcription regulation. Finally, we also observed heart, brain, epidermal growth factor development and nervous system development GO terms enriched for Dyn RT states, which is consistent with the fact that the Dyn RT states are the RT states which change between different cell types.

**Table 1:** RT state analysis on Biological Process GO terms. The colour of the table green and red represent respectively the GO term enriched in Early or Late state. The table highlight only some results the global results are in the Supplementary Table 5.

| GO term | ID | number of genes | Average RT score | Std dev RT score |
| --- | --- | --- | --- | --- |
| translational initiation | GO:0006413 | 123 | 2.683 | 2.188 |
| regulation of mRNA stability | GO:0043488 | 106 | 2.953 | 2.424 |
| rRNA processing | GO:0006364 | 188 | 2.979 | 2.379 |
| mRNA splicing, via spliceosome | GO:0000398 | 215 | 3.005 | 2.695 |
| tumor necrosis factor-mediated signaling pathway | GO:0033209 | 118 | 3.186 | 2.749 |
| regulation of complement activation | GO:0030449 | 104 | 8.260 | 3.739 |
| complement activation, classical pathway | GO:0006958 | 137 | 9.139 | 3.204 |
| detection of chemical stimulus involved in sensory perception of smell | GO:0050911 | 391 | 9.486 | 3.664 |
| sensory perception of smell | GO:0007608 | 226 | 9.535 | 4.019 |

**Table 2:** RT state analysis Molecular Function GO terms. The colour of the table green and red represent respectively the GO term enriched in Early or Late state. The table highlight only some results the global results are in the Supplementary Table 5.

| GO term | ID | Number of genes | Average RT score | Std dev RT score |
| --- | --- | --- | --- | --- |
| structural constituent of ribosome | GO:0003735 | 159 | 3.157 | 2.608 |
| protein N-terminus binding | GO:0047485 | 100 | 3.270 | 2.996 |
| unfolded protein binding | GO:0051082 | 115 | 3.313 | 2.645 |
| RNA binding | GO:0003723 | 1267 | 3.317 | 2.865 |
| mRNA binding | GO:0003729 | 112 | 3.321 | 3.062 |
| G-protein coupled receptor activity | GO:0004930 | 638 | 7.937 | 4.163 |
| olfactory receptor activity | GO:0004984 | 391 | 9.486 | 3.664 |
| immunoglobulin receptor binding | GO:0034987 | 72 | 9.500 | 3.014 |
| antigen binding | GO:0003823 | 158 | 9.513 | 2.987 |
| odorant binding | GO:0005549 | 92 | 11.022 | 3.508 |

### Representation of sub-compartment domains and LAD in the RT states

Genome-wide chromatin conformation capture (Hi-C) experiments (32) have revealed Topologically Associated Domains (TADs) (4-6). The majority of these domains were conserved across cell types, but the inter-TAD interactions are found to be cell type-specific (4, 32, 36). Comparing TAD and RT states allows us to analyze changes in the 3D chromatin organization that occur during cell differentiation which respect to RT states. Furthermore in higher organization of the genome was identified, A and B compartments characterized by the spatial segregation of open and closed chromatin (32).

Rao and collaborators analyzed the A and B compartments and detected six subcompartments based on their contact, architecture and epigenetics features: A1 and A2 (part of the A compartment) and B1, B2, B3 and B4 (part of the B compartment). In this analysis, we ignored the B4 compartment, which is defined only for by a handful of regions, containing many KRAB-ZNF superfamily genes and 0.01% of the bins of each RT possess overlapped with this sub-compartment (33). The A1 sub-compartment is characterised by a high density of actively expressed genes, and enriched with permissive histone marks (H3K36me3, H3K79me2, H3K27ac, and H3K4me1 and H3K4me3) are associated with active transcription (33). We found the A1 sub-compartment highly enriched in Early1 and Early2 RT states (85% and 62% of bins in these two states belong to this sub-compartment respectively) and the proportion reduced steadily to almost zero for the B2 and B3 sub-comportments. In addition of the afore mentioned euchromatin marks, the A2 sub-compartment is also associated with the repressive mark H3K9me3, and is enriched in the states Early3 (59% of the bins) and Dyn1 to Dyn4 (Dyn1: 39%, Dyn2: 56%, Dyn3: 45%) (Fig 6A) and depleted in all others. These five RT states are mainly dynamics and the genes located within these RT states are differentially expressed among different the cell lines analyzed (Fig 5B), so it is expected to have these different states potentially associated with repressive chromatin marks.

The RT states enriched at A1 and A2 sub-compartments are also depleted in LADs (Fig 6A and 6B), which is consistent with the fact that LADs are made of facultative heterochromatin with late replicating properties and correlated positively with H3K27me3 and negatively with H3K36me3. In contrast, sub-compartment B1 with characteristics of facultative heterochromatin (is found to be enriched in Dyn1 to Dyn6 RT states (Dyn1: 31%, Dyn2: 20%, Dyn3: 23%, Dyn4: 30%, Dyn5: 14% and Dyn6: 25%) (Fig 6A). The sub-compartments B2 and B3, which correspond to regions that are not replicated until the end of S phase, show the opposite pattern of sub-compartment A1, which is enriched in late RT states including Dyn9 (27% and 64% respectively), Late1 to Late3 (Late1: 34% and 59%, Late2: 32% and 65%, and Late3: 27% and 71%,respectively) (33) and depleted in early RT states (Fig 6A).

LADs are formed by the interaction between nuclear lamina and the genome and are involved in genome organisation. Most of the LADs are located in gene poor, A/T rich regions, and regions enriched in repetitive elements such as SINEs and LINEs (8). The RT states enriched in B2, and B3 sub-compartments are also enriched in LADs. As an example, around 60% of bins in Late RT states (Dyn9, Late1-3) contain LADs. In contrast, only approximately 2.5% of bins in Early1 RT state overlap with LADs (Fig 6B). Therefore, the Late RT states possess all the characteristics of condensed heterochromatin, late replication and dense in LAD domains. Some of the LADs are cell-type specific and defined as facultative LADs (fLADs), and others are conserved across different cell types and are referred to as constitutive LADs (cLADs) (8). We speculate the subset of LADs with Late RT states (Dyn9, Late1-3) that are likely to be cLADs, and are enriched in B3 sub-compartment (Fig 6A and B) and A/T rich, gene poor regions (Fig 4 and Supplementary Figure 6). The fLADs, which are cell specific, on the other hand, could be localized more in the Dyn RT states, and enriched in B1 and B2 sub-compartment, which are correlated with the facultative heterochromatin.

## DISCUSSION

Genome-wide datasets constitute a valuable tool for genome annotation of distinct chromatin functional properties, especially the non-coding part of the genome. Histone modification, DNA methylation, chromatin accessibility and disease association have all been used to provide functional annotation (1, 37-41). Since RT is a fundamental biological process, closely related to chromatin organization and transcription regulation dynamics during cell fate commitment, we believe RT can supplement the current functional annotation of the genome. Our RT state concept is inspired by the chromHMM work of Ernst and Kellis (9-11). They define chromatin states by applying HMM on ChIP-seq data of histone modifications, together with histone variants (H2A.Z), Polymerase II and CTCF. The chromatin state defined possesses specific functional, experimental, conservation, annotation, and sequence-motif enrichments.

In this work, we develop a novel Hidden Markov Model to classify genome-wide RT programs from 42 datasets into 15 RT states such that these RT states can be used to annotate the genome. The RT state annotation improves over the RD annotation (constitutive and developmental) introduced earlier (24, 42) by offering richer and more quantitative annotation with better resolution. Although both the RT and the chromatin states are developed from applying HMM to demarcate the chromosomes into segments of diverse flavours, the RT and chromatin states have different properties and are complementary to each other. In addition to the source of information, the major differences between RT and chromatin states are four-fold: 1) while chromatin states change from one cell type to another, RT states is not cell type specific. This is because RT states are summarized from a compendium of cell types and developmental stages. 2) the chromatin states are categorical whereas the RT values are ordinal and quantitative, smaller RT state values indicates earlier replication; 3) chromatin state is fine grain type annotation, the resolution is typically 200bp. RT states are defined on a much larger scale, annotated at 50kb level throughout the genome as RT status typically remain stable over a long stretch of the chromosome in most cell types; 4) the abundance of the 15 chromatin states vary dramatically while the 15 RT states have similar proportions across the genome.

The RT states defined by the HMM method illustrate functional properties of the genome. Early RT states possessed open chromatin features with high gene density, enriched in A1 and A2 sub-compartments, enriched in epigenetic marks associated with open chromatin (H3K36me3, H3K79me2, H3K27ac, H3K4me1) and depleted in LADs. Late RT states possessed heterochromatin properties, rich in pseudogenes and lncRNAs, weak in genes and enriched in LADs and B2 and B3 sub-compartments. Dynamic RT states include chromosomal regions that change RT during development (22, 24) and were linked to more variable gene expression activity, chromatin organization, GO terms relative to the development of tissues and possess the characteristics of facultative heterochromatin. The novel RT state concept we propose here has many applications. For example, an application could be to calculate the average RT scores for a set of genomic entities such as genes belong to a particular gene set of pathway. This is possible because the RT states are ordinal (low RT state number indicates early RT and high RT state number indicates late RT). An unusually low or high average RT scores suggest RT connection among these entities. We can also compare the RT scores between two groups of genomic entities of interest to see if they share the same RT profile.

In addition to the reductionist style summary of multiple series of RT values from different cell types, RT states also offer easy visualization of RT data as shown in the wave-like pattern observed in Fig 3C, suggesting early or late replicating genomic regions are arranged in an ordered fashion. Comparing to plots of individual RT values shown at the bottom of the Fig 3C, the pattern of RT states is rather clean and smoother, and a single track of RT states is sufficient to give researchers an overview of the local RT landscape as oppose to 42, highlighting the benefit of the model-based approach we took to obtain RT states. As shown in Fig 6A, the RT states are closely related to the A/B compartment concept derived from Hi-C data, early RT states match compartment A while late states correspond to compartment B. This also corroborate well with the computational derived 3D chromosomal organization constructed based on Hi-C data (Fig 3F in Hu et al. 2013 (43)), confirming the intimate relationship among RT, A/B compartments in the chromosomes and 3D shapes of the chromosomes (44).

Various computational methods have been proposed to identify RDs using replication timing profile data (16, 26, 44). In a recent study, Liu et al. described an advanced deep neural network (DNN)-based method named DNN-HMM that is capable of identifying multiple RDs genome-wide (22). Using manually labelled training data, Liu and colleagues apply DNN on Repliseq data to assign four major types (and 14 subtypes) of RDs throughout the genome. On top of the methodology, the biggest difference between our method and DNN-HMM (and other existing RD calling methods) is that DNN-HMM is designed to call RDs in a single cell type using replication profiling data from that cell type whereas we are trying to define and call RT states across a compendium of cell types, emphasizing the homogeneity and heterogeneity of the RT programs among cells from different developmental lineages and stages. Each of the 15 RT states represent a particular combination of RDs from multiple cell types. As a results, the 15 RT states can be used to annotate the diverse properties of the genome holistically, not just for a single cell type. We believe our method represents the first attempt to jointly model multiple RT profiling data in order to reveal underlying biological insights.

Many studies have been conducted using epigenetics to annotate the genome, especially the non-coding part, RT states can add a new dimension to the annotation, and can elucidate the changes in organisation during differentiation and development. This work could also shed light on the dynamics of the 3D chromatin organisation during differentiation by correlating the RT states and the Hi-C data. Data with better resolution (smaller size of fragments) of the RT data will allow for better definition of the domains of replication and refinement of our models. This could permit future analysis of the RT of specific promoters involved in cell differentiation.

## MATERIAL AND METHODS

### Replication timing data

We analysed 42 genome-wide RT datasets of Repli-chip previously published (24), representing 25 cell types and differentiation intermediates of human development. The genome was first divided into bins of 50kb, then RT value was calculated for each bin. The 42 datasets include embryonic stem cells (hESC), primary cells and established cell lines representing intermediate stages of endoderm, mesoderm, ectoderm, and neural crest (NC) development (Supplementary table 1) (6, 17, 24, 45, 46). Telomeric and centromeric regions, as well as sex chromosomes were removed in this analysis. The RT values used in our analysis cover 85% of the human genome. The distributions of the RT values aggregated across all samples are shown in Fig 1A. RT data were normalized using the limma package (47) in R and rescaled to the same range using quantile normalization (48).

### Hidden Markov Model

Like the chromHMM model proposed by Ernst and Kellis to call chromatin states (9-11), we use a multivariate HMM to capture the underlying structure of the 42 RT programs. Our model assumes a fixed number--15, of hidden states *K* (the selection of *K* is discussed in detail in a subsequent section). For each hidden state, we use a distinct normal distribution (emission probabilities) to characterize the RT values (at the bin level) from each RT program. Specifically, for each of the *K* hidden states, there are two parameters: *µ*_*k*_ and 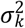 denoting the mean and variance of the normal distribution in hidden state *k* _(*k*= 1, …,*K*)_ that the RT values follow. Although many of the datasets and cell types are closely related, for the sake of mathematical simplicity, following chromHMM, we treat the RT values obtained from different cell types as independent condition on the assigned RT state at the locus. For a specific chromosome *C*, The full likelihood of all of the observed RT values *x* and together with key HMM parameters *a* (initial probabilities), *b* (transition probabilities) can be expressed:

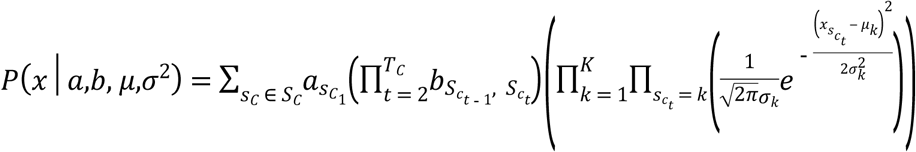

Where *N*(,) is the density function for normal distribution *c_t_* denotes a 50kb bin on chromosomes where *t* = 1,…*Tc*. Let *b*_*i,j*_denotes the probability of transitioning from state *i* to *j* where *i*_(*i*=1,…,*K*)_ and *j*_(*j*=1,…,*K*)_. We also have parameters *a*_*i* (*i*=1,…,*K*)_ which denote the probability that the state of the first interval on the chromosome is *i*. Let *S*_*c*_ ∈ *S*_*c*_ be an unobserved state sequence through chromosome _*c*_ and *S*_*c*_ be the set of all possible state sequences. Let 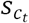 denote the unobserved state on chromosome *c* at location *t* for state sequence *S*_*c*_. The model has a similar structure as chromHMM method (9) but customized for fitting RT data. We use R package depmix4 to conduct inference on the HMM (49). Once the HMM parameters are estimated and 50 kb bins along the genomes are assigned to different RT states. We group all bins belong to the same RT states together and obtain average RT values for each RT state in every cell type. We next perform a two-way clustering on these average RT values, and ordered the 15 RT states such that the first one has the earliest RT values and the last one has the latest RT values. By doing this, the 15 RT states are ordered from early RT to late RT.

### Annotation of RT states

#### Compare to chromatin states

Chromatin state annotations of nine ENCODE cell types (9) defined on nine different epigenetic marks were downloaded from the chromHMM website (http://compbio.mit.edu/ChromHMM/). The nine cell lines are: GM12878 (B-lymphocyte), H1-hESC (human embryonic stem cells), HepG2 (hepatocellular carcinoma), HMEC (mammary epithelial cells), HSMM (skeletal muscle myoblasts), K562 (leukemia), NHEK (epidermal keratinocytes), NHLF (lung fibroblasts) and HUVEC (umbilical vein endothelial cells). The chromHMM 15 chromatin states are: active promoter, weak promoter, inactive/poised promoter, strong enhancer, strong enhancer, weak/poised enhancer, weak/poised enhancer, insulator, transcriptional transition, transcriptional elongation, weak transcribed, Polycomb-repressed, heterochromatin, low signal, repetitive/Copy Number Variation 1 and repetitive/Copy Number Variation 2. The 15 chromatin states were defined based on the occurrence of a combination of eight different epigenetic marks (H3K27ac, H3K27me3, H3k36me3, H3K4me1, H3K4me2, H3K4me3, H3K9ac and H3K20me1) and CTCF binding sites. To compare RT states and chromatin states, the proportion of the genomes corresponds to the 15 chromatin states and the 15 RT states were obtained with the following formula: 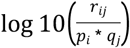. With chromatin state proportions: *p*_(*p*1, …, *p*15)_; and RT state proportions: *q*_(*q*1, …,*q*15)_, *r*_*i*(*i*= 1,.., 15)_, *j*_(*j*= 1,…,15)_ corresponds to the combination of chromatin states *i* and RT state *j*.

#### Gene features analysis

The detection of the gene features in each 50 kb bin was performed on Hg19 General Feature Format file (file available on www.gencodegenes.org). There are four different global gene features: protein coding (contains an open reading frame), ncRNA (non-coding RNA), lncRNA (long non-coding RNA) and pseudogenes (similar to known proteins but contain a frameshift and/or stop codon, which disrupts the open reading frame).

#### Transcriptome analysis on protein coding genes

Expression level measured by the Reads Per Kilobase of transcript per Million mapped reads (RPKM) of RNA sequencing data of protein coding genes of 17 cell lines were downloaded from REMC (http://www.roadmapepigenomics.org/) (50). The 17 cell types include the nine ENCODE cell types used in the chromHMM analysis plus additional eight cell types overlapping with the 42 cell lines used to construct the RT states: H1 BMP4 derived trophoblast, H1 derived mesenchymal stem, H1 derived neuronal progenitor, hESC derived CD184 endoderm, hESC derived CD56 ectoderm, hESC derived CD56 mesoderm, HELA. The RPKM value of the protein coding genes that located inside regions corresponding to each of the 15 RT states were calculated, then their average expression levels in is computed in log scale as log10(RPKM_average_).

#### Gene ontology term enrichment analysis

Gene ontology (GO) (51) term enrichment analysis is performed using Proteinside (52). The analysis is realised using the lists of protein coding genes belong to each of the 15 RT states compiled for the RNA-seq analysis. The twenty GO terms most enriched in each of the 15 RT states were used to analyse the specificity and characteristics of genes localized in each RT state. We also computed the average RT score for all the GO terms in the Biological Process (BP) and Molecular Function (MF) categories (51). Since the RT states are ordered from Early to Late, the average RT score for a group of genes reflects their replication timing.

#### Three-dimensional chromosomal organization analysis

We use high resolution Hi-C data (GEO accession number: GSE63525) obtained from human lymphoblastoid cells (GM12878) produced by Rao and collaborators (33). We count the number of contact domains found in regions corresponding to each of the 15 RT states. The same method was used to analyse the Lamin Associated Domains (LADs) in fibroblasts. The LADs data was obtained from ENCODE (7).

## AVAILABILITY

R scripts for inferring HMM models and Perl scripts for further analysis are available https://github.com/PouletAxel/script_HMM_Replication_timing

## SUPPLEMENTARY DATA

Supplementary file: Supplementary figure.doc, Supplementary table5_GOterm.xls and HMM_RT_hg19_srt.bed

## ACKNOWLEDGEMENT

We thank the members of the Gilbert Lab and the Corces Lab for sharing the RT data and for discussions. We thank Professor Christophe Tatout for comments on the draft of the manuscript. And we thank Michael Holden Nichols in the Corces lab for comments on the draft of the manuscript.

## REFERENCES

1. Dunham I, Kundaje A, Aldred SF, Collins PJ, Davis C, Doyle F, et al. An integrated encyclopedia of DNA elements in the human genome. Nature. 2012;489(7414):57–74.

2. Bernstein BE, Stamatoyannopoulos JA, Costello JF, Ren B, Milosavljevic A, Meissner A, et al. The NIH Roadmap Epigenomics Mapping Consortium. Nat Biotechnol. 2010;28(10):1045–8.

3. Bae JB. Perspectives of international human epigenome consortium. Genomics Inform. 2013;11(1):7–14.

4. Dixon JR, Selvaraj S, Yue F, Kim A, Li Y, Shen Y, et al. Topological domains in mammalian genomes identified by analysis of chromatin interactions. Nature. 2012;485(7398):376–80.

5. Hou CH, Li L, Qin ZHS, Corces VG. Gene Density, Transcription, and Insulators Contribute to the Partition of the Drosophila Genome into Physical Domains. Mol Cell. 2012;48(3):471–84.

6. Sexton T, Yaffe E, Kenigsberg E, Bantignies F, Leblanc B, Hoichman M, et al. Three-Dimensional Folding and Functional Organization Principles of the Drosophila Genome. Cell. 2012;148(3):458–72.

7. Guelen L, Pagie L, Brasset E, Meuleman W, Faza MB, Talhout W, et al. Domain organization of human chromosomes revealed by mapping of nuclear lamina interactions. Nature. 2008;453(7197):948–51.

8. Meuleman W, Peric-Hupkes D, Kind J, Beaudry JB, Pagie L, Kellis M, et al. Constitutive nuclear lamina-genome interactions are highly conserved and associated with A/T-rich sequence. Genome Res. 2013;23(2):270–80.

9. Ernst J, Kheradpour P, Mikkelsen TS, Shoresh N, Ward LD, Epstein CB, et al. Mapping and analysis of chromatin state dynamics in nine human cell types. Nature. 2011;473(7345):43–9.

10. Ernst J, Kellis M. ChromHMM: automating chromatin-state discovery and characterization. Nat Methods. 2012;9(3):215–6.

11. Ernst J, Kellis M. Discovery and characterization of chromatin states for systematic annotation of the human genome. Nat Biotechnol. 2010;28(8):817–25.

12. Hiratani I, Ryba T, Itoh M, Yokochi T, Schwaiger M, Chang CW, et al. Global Reorganization of Replication Domains During Embryonic Stem Cell Differentiation. Plos Biol. 2008;6(10):2220–36.

13. Ryba T, Battaglia D, Pope BD, Hiratani I, Gilbert DM. Genome-scale analysis of replication timing: from bench to bioinformatics. Nat Protoc. 2011;6(6):870–95.

14. Hiratani I, Takebayashi S, Lu J, Gilbert DM. Replication timing and transcriptional control: beyond cause and effect--part II. Curr Opin Genet Dev. 2009;19(2):142–9.

15. Pope BD, Gilbert DM. The replication domain model: regulating replicon firing in the context of large-scale chromosome architecture. J Mol Biol. 2013;425(23):4690–5.

16. Hansen RS, Thomas S, Sandstrom R, Canfield TK, Thurman RE, Weaver M, et al. Sequencing newly replicated DNA reveals widespread plasticity in human replication timing. Proc Natl Acad Sci U S A. 2010;107(1):139–44.

17. Hiratani I, Ryba T, Itoh M, Rathjen J, Kulik M, Papp B, et al. Genome-wide dynamics of replication timing revealed by in vitro models of mouse embryogenesis. Genome Res. 2010;20(2):155–69.

18. Gerhardt J, Zaninovic N, Zhan Q, Madireddy A, Nolin SL, Ersalesi N, et al. Cis-acting DNA sequence at a replication origin promotes repeat expansion to fragile X full mutation. J Cell Biol. 2014;206(5):599–607.

19. Rivera-Mulia JC, Desprat R, Trevilla-Garcia C, Cornacchia D, Schwerer H, Sasaki T, et al. DNA replication timing alterations identify common markers between distinct progeroid diseases. Proc Natl Acad Sci U S A. 2017;114(51):E10972–E80.

20. Ryba T, Battaglia D, Chang BH, Shirley JW, Buckley Q, Pope BD, et al. Abnormal developmental control of replication-timing domains in pediatric acute lymphoblastic leukemia. Genome Res. 2012;22(10):1833–44.

21. Sasaki T, Rivera-Mulia JC, Vera D, Zimmerman J, Das S, Padget M, et al. Stability of patient-specific features of altered DNA replication timing in xenografts of primary human acute lymphoblastic leukemia. Exp Hematol. 2017;51:71-82 e3.

22. Liu F, Ren C, Li H, Zhou P, Bo X, Shu W. De novo identification of replication-timing domains in the human genome by deep learning. Bioinformatics. 2016;32(5):641–9.

23. Pope BD, Ryba T, Dileep V, Yue F, Wu WS, Denas O, et al. Topologically associating domains are stable units of replication-timing regulation. Nature. 2014;515(7527):402-+.

24. Rivera-Mulia JC, Buckley Q, Sasaki T, Zimmerman J, Didier RA, Nazor K, et al. Dynamic changes in replication timing and gene expression during lineage specification of human pluripotent stem cells. Genome Res. 2015;25(8):1091–103.

25. Hu M, Deng K, Qin Z, Liu JS. Understanding spatial organizations of chromosomes via statistical analysis of Hi-C data. Quant Biol. 2013;1(2):156–74.

26. Pope BD, Ryba T, Dileep V, Yue F, Wu W, Denas O, et al. Topologically associating domains are stable units of replication-timing regulation. Nature. 2014;515(7527):402–5.

27. Rivera-Mulia JC, Gilbert DM. Replicating Large Genomes: Divide and Conquer. Mol Cell. 2016;62(5):756–65.

28. Hatton KS, Dhar V, Brown EH, Iqbal MA, Stuart S, Didamo VT, et al. Replication program of active and inactive multigene families in mammalian cells. Mol Cell Biol. 1988;8(5):2149–58.

29. Marchal CS, T.; Korey Wilson, V.; Sima, J.; Rivera-Mulia, J.; Trevilla Garcia, C.; Nogues C.; Nafie, E.; Gilbert D. M. Repli-seq: genome-wide analysis of replication timing by next-generation sequencing. bioRxiv. 2017.

30. Naidoo N, Pawitan Y, Soong R, Cooper DN, Ku CS. Human genetics and genomics a decade after the release of the draft sequence of the human genome. Hum Genomics. 2011;5(6):577–622.

31. Gilbert N, Boyle S, Fiegler H, Woodfine K, Carter NP, Bickmore WA. Chromatin architecture of the human genome: gene-rich domains are enriched in open chromatin fibers. Cell. 2004;118(5):555–66.

32. Lieberman-Aiden E, van Berkum NL, Williams L, Imakaev M, Ragoczy T, Telling A, et al. Comprehensive mapping of long-range interactions reveals folding principles of the human genome. Science. 2009;326(5950):289–93.

33. Rao SSP, Huntley MH, Durand NC, Stamenova EK, Bochkov ID, Robinson JT, et al. A 3D Map of the Human Genome at Kilobase Resolution Reveals Principles of Chromatin Looping. Cell. 2014;159(7):1665–80.

34. Gibcus JH, Samejima K, Goloborodko A, Samejima I, Naumova N, Nuebler J, et al. A pathway for mitotic chromosome formation. Science. 2018;359(6376).

35. Koren A, Polak P, Nemesh J, Michaelson JJ, Sebat J, Sunyaev SR, et al. Differential relationship of DNA replication timing to different forms of human mutation and variation. Am J Hum Genet. 2012;91(6):1033–40.

36. Nora EP, Lajoie BR, Schulz EG, Giorgetti L, Okamoto I, Servant N, et al. Spatial partitioning of the regulatory landscape of the X-inactivation centre. Nature. 2012;485(7398):381–5.

37. Boyle AP, Hong EL, Hariharan M, Cheng Y, Schaub MA, Kasowski M, et al. Annotation of functional variation in personal genomes using RegulomeDB. Genome Res. 2012;22(9):1790–7.

38. Fu Y, Liu Z, Lou S, Bedford J, Mu XJ, Yip KY, et al. FunSeq2: a framework for prioritizing noncoding regulatory variants in cancer. Genome Biol. 2014;15(10):480.

39. Ward LD, Kellis M. HaploReg: a resource for exploring chromatin states, conservation, and regulatory motif alterations within sets of genetically linked variants. Nucleic Acids Res. 2012;40(Database issue):D930–4.

40. Ward LD, Kellis M. HaploReg v4: systematic mining of putative causal variants, cell types, regulators and target genes for human complex traits and disease. Nucleic Acids Res. 2016;44(D1):D877–81.

41. Chen L, Qin ZS. traseR: an R package for performing trait-associated SNP enrichment analysis in genomic intervals. Bioinformatics. 2016;32(8):1214–6.

42. Dileep V, Rivera-Mulia JC, Sima J, Gilbert DM. Large-Scale Chromatin Structure-Function Relationships during the Cell Cycle and Development: Insights from Replication Timing. Cold Spring Harb Symp Quant Biol. 2015;80:53–63.

43. Hu M, Deng K, Qin Z, Dixon J, Selvaraj S, Fang J, et al. Bayesian inference of spatial organizations of chromosomes. PLoS Comput Biol. 2013;9(1):e1002893.

44. Ryba T, Hiratani I, Lu J, Itoh M, Kulik M, Zhang J, et al. Evolutionarily conserved replication timing profiles predict long-range chromatin interactions and distinguish closely related cell types. Genome Res. 2010;20(6):761–70.

45. Pope BD, Tsumagari K, Battaglia D, Ryba T, Hiratani I, Ehrlich M, et al. DNA Replication Timing Is Maintained Genome-Wide in Primary Human Myoblasts Independent of D4Z4 Contraction in FSH Muscular Dystrophy. Plos One. 2011;6(11).

46. Ryba T, Hiratani I, Sasaki T, Battaglia D, Kulik M, Zhang J, et al. Replication timing: a fingerprint for cell identity and pluripotency. PLoS Comput Biol. 2011;7(10):e1002225.

47. Smyth GK. limma: linear models for microarray data In Bioinformatics and computational biology solutions using R and bioconductor. ed. Gentleman R ea, editor. New York 2005. pp. 397–420 p.

48. Irizarry RA, Hobbs B, Collin F, Beazer-Barclay YD, Antonellis KJ, Scherf U, et al. Exploration, normalization, and summaries of high density oligonucleotide array probe level data. Biostatistics (Oxford, England). 2003;4(2):249–64.

49. Visser IS, M. depmixS 4: An R Package for Hidden Markov Models. Journal of Statistical Software. 2010;36:1–21.

50. Bernstein BE, Stamatoyannopoulos JA, Costello JF, Ren B, Milosavljevic A, Meissner A, et al. The NIH Roadmap Epigenomics Mapping Consortium. Nat Biotechnol. 2010;28(10):1045–8.

51. Ashburner M, Ball CA, Blake JA, Botstein D, Butler H, Cherry JM, et al. Gene ontology: tool for the unification of biology. The Gene Ontology Consortium. Nat Genet. 2000;25(1):25–9.

52. Kaspric N, Reichstadt M, Picard B, Tournayre J, Bonnet M. Protein Function Easily Investigated by Genomics Data Mining Using the ProteINSIDE Online Tool Genomics and Computational Biology 2015;Vol. 1.

